# Forecasting multi-trait resistance evolution under antibiotic stress

**DOI:** 10.1101/2025.10.17.683023

**Authors:** Suvam Roy, Eric Libby, Peter Lind

## Abstract

Many bacteria rely on efflux pumps to survive antibiotic stress, and exposure to antibiotics often leads to mutations in pump genes or their regulators that increase pump expression. Predicting the spectrum of these mutations is important for designing effective antibiotic treatments, but the underlying regulatory networks are large and complex, making them difficult to map experimentally. To address this challenge, we developed a mathematical framework that integrates dynamical equations for efflux pump regulation with a genetic algorithm for parameter estimation and evolutionary simulations. Using this framework, we simulated *in silico* evolution of *Pseudomonas aeruginosa* under exposure to the antibiotics meropenem, tobramycin, and ciprofloxacin. The simulations revealed mutational spectra affecting the expression of four RND efflux pumps and their shared regulatory network. The most frequently mutated genes were single-target regulators that matched well with previous observations in clinical and *in vitro* studies. The model also showed that the shared use of the OprM protein by two pumps is a key factor shaping their distinct mutational patterns. Mutations often produced multi-trait phenotypes, manifesting as collateral sensitivity or cross-resistance to antibiotics not used for selection. While cross-resistance evolved readily, its extent depended on initial pump expression levels and thus may vary between strains. Finally, simulations of changing environments showed that efflux pump genes tend to be lost in the absence of antibiotics, suggesting a potential strategy to steer bacterial evolution toward reduced capacity to re-evolve resistance.

## 1 Introduction

Bacteria regularly encounter toxic compounds, such as antibiotics, that threaten their survival. A common defence mechanism involves the use of efflux pumps that expel diverse toxic compounds from inside the cell [1]. Indeed, when exposed to antibiotics, bacteria often evolve multidrug resistance by acquiring mutations that cause overexpression of these genes [2, 3]. Interestingly, the regulation of efflux pump systems is remarkably complex [4, 5]. Their expression is controlled by intricate hierarchical networks of both local and global transcription factors (TFs) as well as post-translational and post-transcriptional regulation [5, 6]. A consequence of this complex regulation is that resistance mutations typically do not occur in the efflux pump genes themselves but rather in regulatory genes. Predicting these mutational targets a priori and their functional consequences, e.g., fitness [7] or epistatic effects [8], is complicated by the complexity of interacting components. Moreover, once a mutation has fixed it can alter future mutational targets, having downstream effects on evolutionary trajectories. These effects can lead to medically relevant phenomena such as cross-resistance and collateral sensitivity [9]. Thus, predicting the effects of mutations on efflux pump regulation can have important implications for designing antibiotic treatments to pathogenic bacteria.

One way to develop and test such predictions is to carry out *in vitro* evolution experiments. The problem is that experimentally mapping this mutational landscape across entire regulatory networks is resource intensive [10] due to the vast combinatorial space of possible mutations [11], the need for precise genetic manipulation [12], and the requirement for high-throughput phenotypic screening under dynamic conditions. Consequently, computational and theoretical approaches coupled with evolutionary simulations [13–22], can make important contributions. These theoretical frameworks have been shown to be useful in predicting mutational targets [23] and accounting for how the evolutionary history of a strain, i.e. previous mutations in regulators, can alter future resistance evolution. They are also highly adaptable to changes in gene content in different strains or changes in gene expression in different environments. Importantly, they can be used to search a large combinatorial space of mutations and simulate resistance evolution during sequential [24] or combination [25] exposure to antibiotics, and thereby provide experimentally-testable null models and possible targets for novel therapeutic strategies.

Developing a predictive theoretical approach requires a significant amount of empirical data. A single bacterial species can contain multiple efflux pump systems, each with their own regulators and specificity concerning which antibiotics they export. Since, the different efflux pumps also have cross-regulation, changes in the regulation of one pump may affect another. The requirement for empirical data to effectively model gene regulatory networks highlights the need for a well-characterized model system. To this end, *Pseudomonas aeruginosa* is a good candidate because it is a clinically significant opportunistic pathogen that has been studied extensively. *Pseudomonas aeruginosa* possesses four major ResistanceNodulation-Division (RND)-type efflux pumps linked to antibiotic resistance in the clinic: MexAB-OprM, MexXY-OprM, MexCD-OprJ, and MexEF-OprN [5, 26]. Each pump comprises three components: two Mex proteins (localized to the inner membrane and periplasm) and one Opr protein (embedded in the outer membrane). An overview of the regulators of each pump and the antibiotics extruded by each pump is provided in Table1. During antibiotic treatment *Pseudomonas aeruginosa* commonly evolves multidrug resistance through mutations in regulators, e.g. *nfxB* and *mexZ*, that lead to increased production of efflux pumps. Because of their role in resistance, many of the regulators have been subject to decades of experimental studies, providing a detailed mechanistic understanding of their functions. *Pseudomonas aeruginosa* therefore is an ideal model system to provide fundamental insights into the evolution of gene regulatory networks and phenomena such as cross-resistance and collateral sensitivity [2].

**Table 1.**
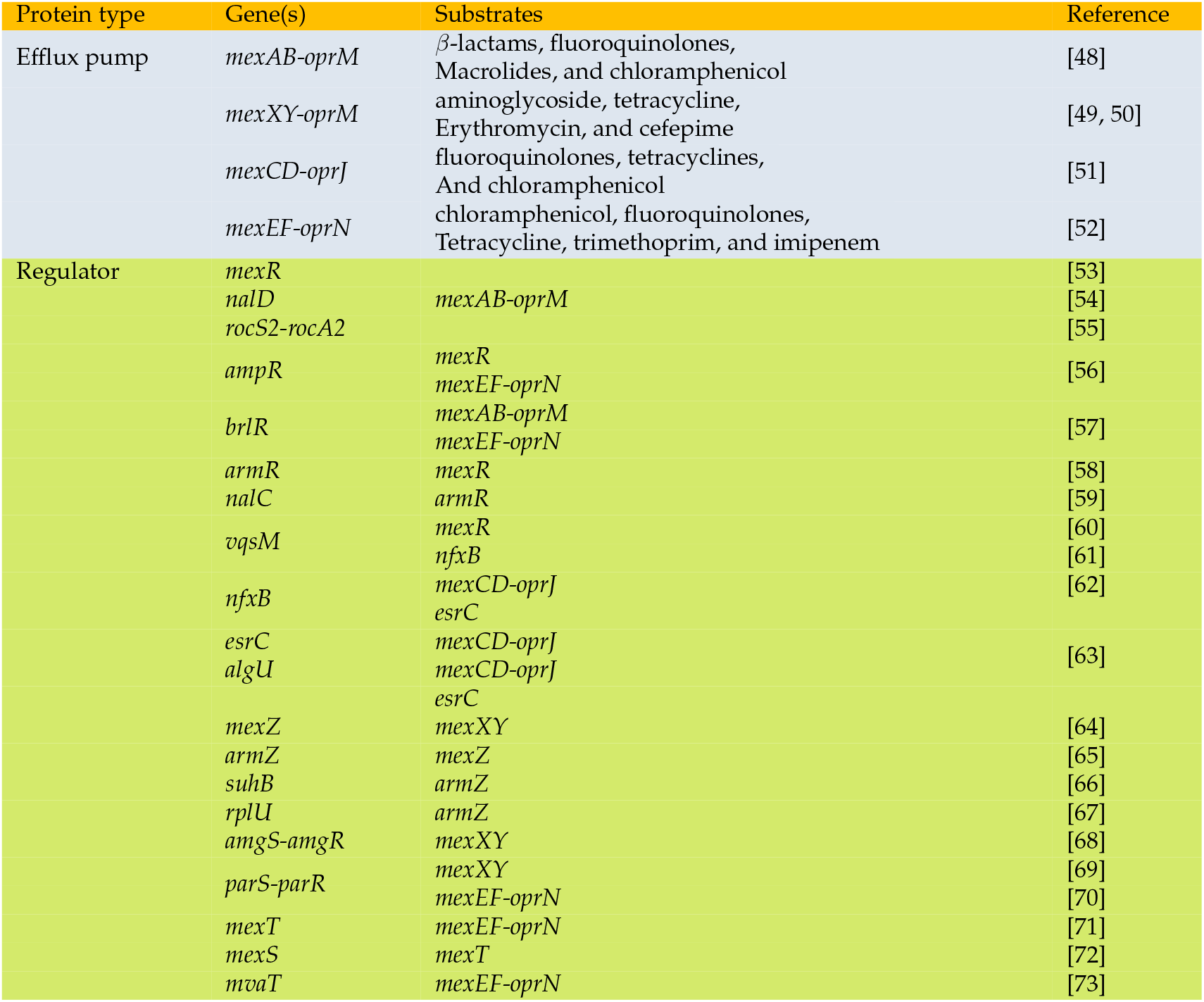
The genes related to the 4 RND efflux pumps in *Pseudomonas aeruginosa* along with their targets (antibiotics or other genes).

Here, we modelled gene expression levels of *Pseudomonas aeruginosa* efflux systems and their regulators following established kinetic modelling techniques [27]. We developed a genetic algorithm to estimate the rates of regulation and simulated the evolutionary trajectories under different antibiotic treatments. From our simulations we identified commonly mutated genes and characterized the diversity of mutational pathways for three clinically relevant antibiotics from different classes (ciprofloxacin, meropenem and tobramycin). We found that the presence of shared regulators and efflux pump components result in diverse patterns of cross-resistance and collateral sensitivity. Upon removal of antibiotics, we observed that resistant mutants acquired deleterious mutations in efflux pump genes as they were costly to produce and no longer useful. Our results provide a detailed analysis of how the complex regulatory networks associated with efflux pumps such as the one of *Pseudomonas aeruginosa* might evolve under antibiotic treatments and produce experimentally testable predictions that might ultimately be useful for guiding clinical treatment regimes.

## 2 Results

### 2.1 Model overview

The interactions in the regulatory networks for four efflux pumps in *Pseudomonas aeruginosa* were defined based on experimental findings as described in Table-1 and shown in Figure-1(a). We described these regulatory interactions using a mathematical model composed of differential equations (see Methods 4.1 for details). In our model the rates of transcription and translation were estimated from experimental data and the two processes were combined into a single-step. The rates in our model pertaining to regulation however, are not known and difficult to measure experimentally. We therefore employed a genetic algorithm to estimate these rates such that the protein levels of the 4 efflux genes reached specifically chosen values. We considered two cases for the protein levels of the efflux genes. In the first case, we fixed the level of all efflux pump proteins to the average protein concentration (200) per cell per gene for *Pseudomonas aeruginosa*. In the second case, we fixed the concentration of MexAB pump to 200, with the concentration of the other three pumps fixed to 40 per cell (as MexAB has been observed to be expressed at a higher rate) [28]. Since the genetic algorithm enacts a stochastic search process, different runs will result in different values for the rates of regulation. Thus, we repeated the fitting process 25 times for each case to study a collection of possible models. The diversity of models and the effects of constraining the efflux protein levels can be found in Figure-S1 and Supplementary section: Robustness of rate estimates.

**Figure 1.**
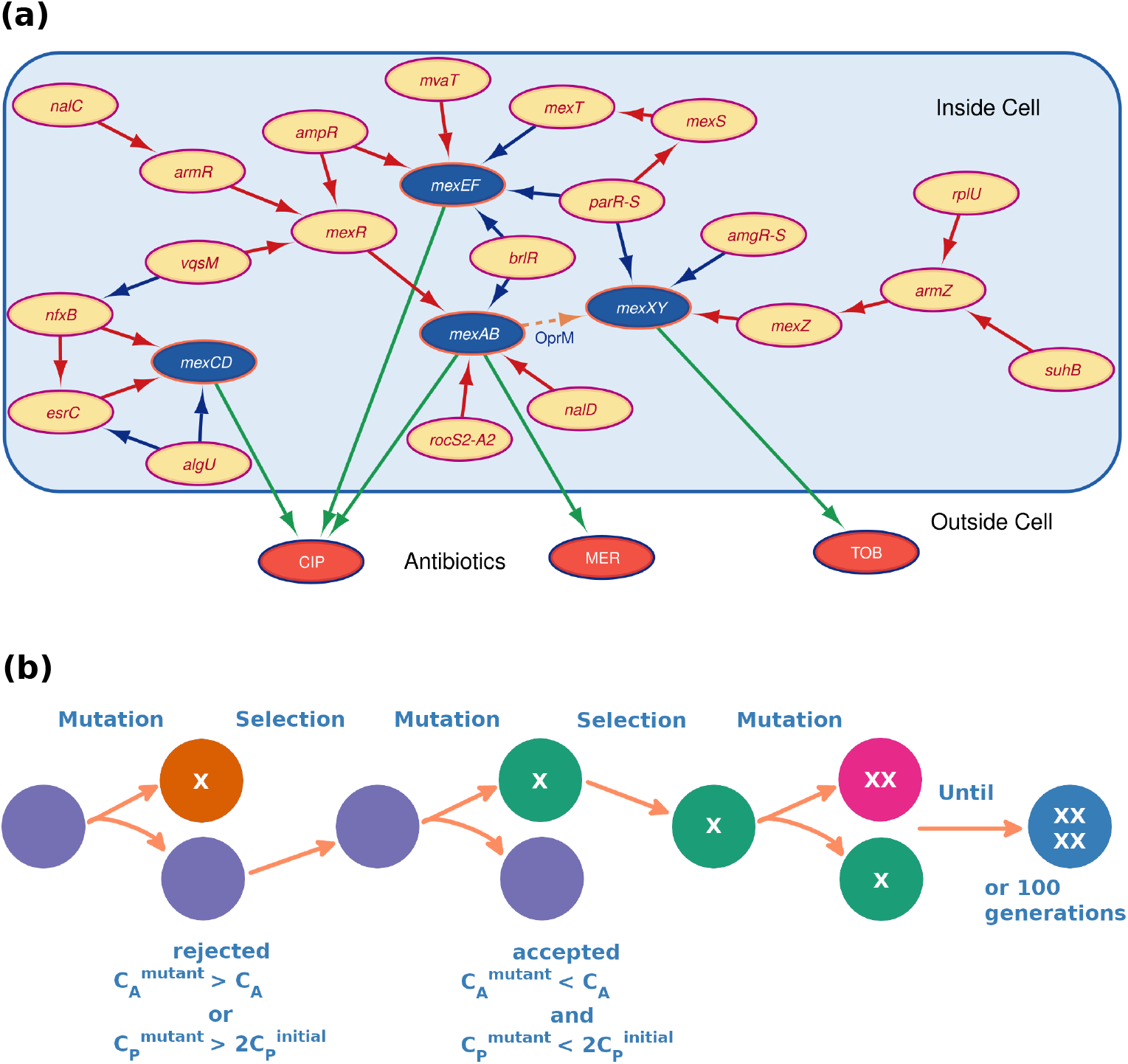
The regulation of efflux pumps and the evolutionary response to antibiotics. **(a):** The diagram shows the gene regulatory network of four major RND efflux pumps in *Pseudomonas aeruginosa*. Blue nodes represent the efflux pump genes: MexAB (*mexA, mexB, oprM*), MexXY (*mexX, mexY, oprM*), MexCD (*mexC, mexD, oprJ*) and MexEF (*mexE, mexF, oprN*). Red nodes represent the three antibiotics we consider: ciprofloxacin (CIP), meropenem (MER), and tobramycin (TOB); and green arrows indicate which antibiotics each pump acts on. Yellow nodes represent regulatory genes with effects indicated by colored arrows: blue is up-regulation and red is down-regulation. The arrow labelled OprM is dashed to indicate that the MexXY pump requires OprM to be functional, and the *oprM* gene is transcribed together with *mexAB* in the *mexAB-oprM* operon. The regulatory proteins RocS2-A2 (RocS2-RocA2), AmgR-S (AmgR-AmS), ParR-S (ParR-ParS) are two-component systems but in our model we consider them as single proteins. **(b):** The schematic shows how evolution acts in our modeling framework. We start with a specific set of parameters describing the dynamics of the efflux regulatory network. Mutations occur randomly in regulatory genes and their benefit is assessed based on two criteria. First, they must increase resistance to antibiotics, represented in the model as lowering the cost of antibiotics (*C*_*A*_). Second, they must not increase the total expression of protein (or cost of protein *C*_*P*_) above some initial threshold, 2 times the initial *C*_*P*_ in the model. This process is repeated until a total of 4 mutations are accepted or 100 mutations have been assessed.

Once we have an initial set of models we then simulate their evolution in response to antibiotics. In our framework adaptation occurs via mutations that cause deletions or duplications, which can also be interpreted as loss-of-function and gain-of-function mutations, respectively. For simplicity, we assume that mutations that increase fitness will fix while those that are neutral or impose a fitness cost are lost. The primary determinant of fitness in our system is the internal antibiotic concentration of cells, which means that mutations that allow cells to more effectively pump out antibiotics will improve fitness. In the absence of any other determinant of fitness, mutations leading to increased production of efflux pumps would fix, regardless of whether antibiotics are present. To avoid such biologically unrealistic results, we add a constraint that mutations must not lead to substantial increases in total protein production, thereby enacting a type of energy budget (see Methods 4.2 for details of how antibiotic and protein production cost is calculated). Thus, our simulations represent evolutionary trajectories where each mutation increases the removal of antibiotics while not incurring large increases in protein production. In our model, mutations occur randomly with an equal probability for deletion and duplication events (see Figure-1(b)). Each evolutionary trajectory stops when either 4 mutations have fixed or 100 mutations have been assessed. For each initial model (set of reaction rates) we simulate 25 evolutionary trajectories.

### 2.2 Mutational patterns and diversity

As a result of our evolutionary simulations we can determine the probabilities of deletion and duplication for all regulators in our model (see Figure-2). We first describe mutations that confer resistance to meropenem. Since meropenem is only pumped out by MexAB, the most common mutations affect regulators of MexAB, i.e. deletions of its negative regulators *nalD* and *rocS2-A2* and duplications of its positive regulator *brlR*. Another frequent target is *mexZ* which occurs due to a competitive interaction between the two pumps MexAB and MexXY, the main pump for tobramycin: both pumps need OprM in order to be functional and OprM is expressed along with MexAB. Because MexXY competes with MexAB over OprM, increased MexAB functionality can be achieved by lowering the expression of MexXY, either by duplicating its negative regulator *mexZ* or deleting its positive regulators *parR-S* and *amgR-S*. The deletion of *parR-S* and *amgR-S* is observed only for the case (A) when both pumps have the same initial expression level, because it lowers the expression of MexXY further.

**Figure 2.**
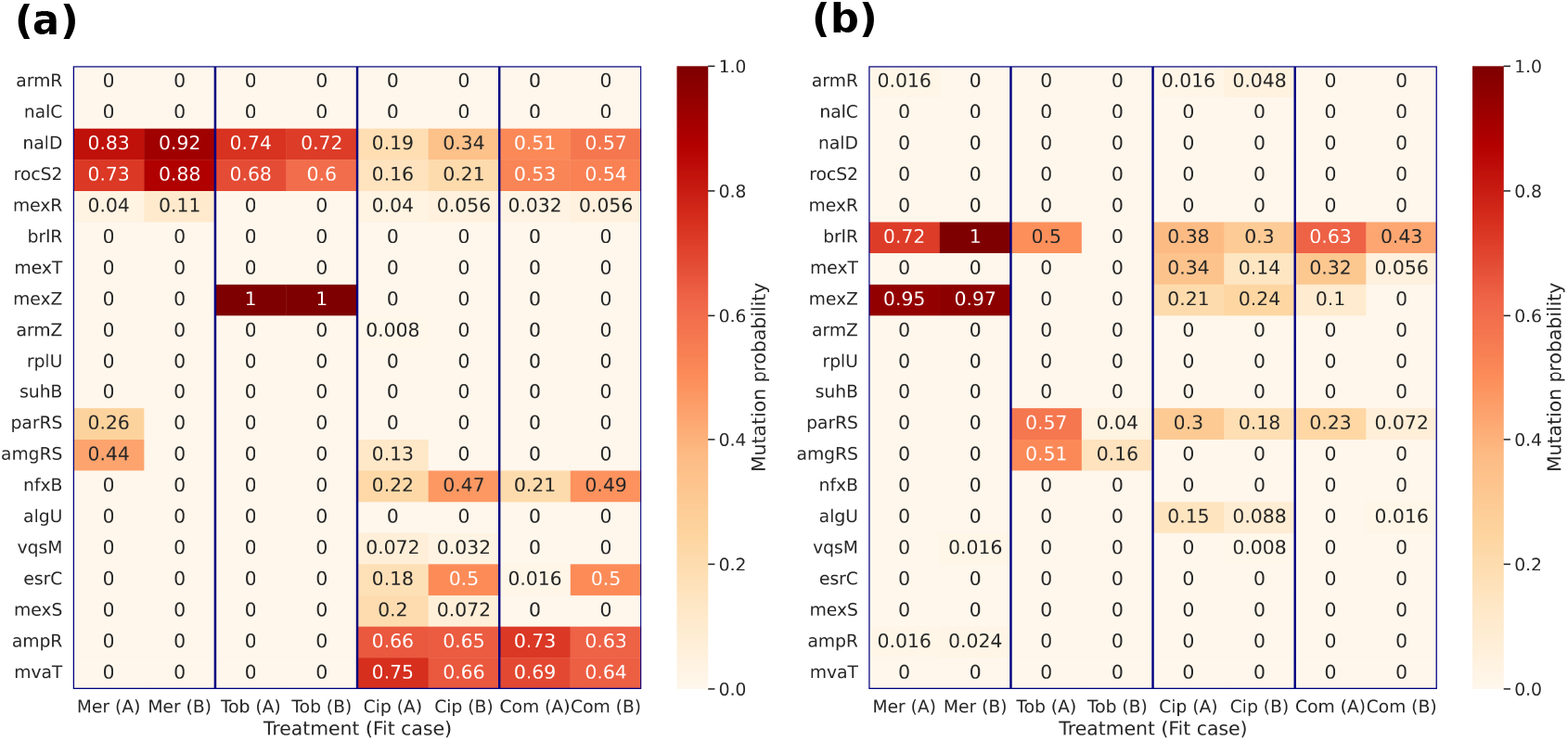
Mutation probabilities of regulatory genes, under different antibiotic treatments. **(a) & (b):** Heatmaps show the probabilities of mutations of different regulatory genes, when evolved in different antibiotic treatments: meropenem (Mer), tobramycin (Tob), ciprofloxacin (Cip), or Com (a combination of all three). The **(a)** heatmap shows deleterious or loss-of-function mutations while the **(a)** heatmap shows gene duplication or gain-of-function mutations. The A and B labels after each antibiotic correspond to the two expression patters considered, i.e. Case A when all four pumps have same expression levels and Case B when the MexAB pump has a higher expression level. The mutational spectra for the two expression cases are largely similar. For both deletions and duplications, regulators with single targets tend to have higher mutation probabilities. The competition between MexAB and MexXY for OprM is an important determinant of the type of mutation. This competition results in deletion of *parRS* and *amgRS* and duplication of *mexZ* under meropenem treatment, while we observed opposite trend under tobramycin treatment.

Since tobramycin is pumped out by MexXY, selection under tobramycin treatment leads to the opposite mutations as in meropenem, i.e. deletions of *mexZ* or duplications of *parR-S* and *amgR-S*. Deletions of *mexZ* occurred in all simulations because it directly leads to increased expression of MexXY. Whether duplications of *parR-S* and *amgR-S* were found depended on the initial steady state concentration of MexXY, i.e. either Case A or Case B. Duplications of *parR-S* and *amgR-S* are more likely to occur in Case A than Case B, because the strength of these two regulators is low in Case B based on their kinetic parameters, thereby reducing their impact. Resistance to tobramycin also occurs via mutational targets *nalD* and *rocS2-A2*, but only after *mexZ* deletions. The reason is that MexXY needs OprM to form a functional efflux pump, and the *oprM* genes is transcribed from the *mexAB-oprM* operon. So increasing the amount of MexAB leads to more OprM and thus more functional MexXY. For the same reason, we found duplications of *brlR* which upregulates MexAB, but only in Case A.

Ciprofloxacin is pumped out by MexAB-OprM, MexCD-OprJ, and MexEF-OprN. Therefore, we found deletions in the negative regulators of these pumps: *nalD* and *rocS2-A2* for MexAB-OprM; *nfxB* and *esrC* for MexCD-OprJ; and *ampR* and *mvaT* for MexEF-OprN. Of these negative regulators, the most common mutational targets were the *mvaT* and *ampR* genes. Both of these regulate MexEF-OprN and do not interfere with the regulation of any other pump, i.e. these nodes are isolated in the regulatory network (see Figure-1). Thus, mutating them can directly lead to increased MexEF-OprM production without any epistatic interactions. In terms of duplications, we found mutations in the positive regulators of pumps: *brlR* (for MexAB-OprM and MexEF-OprN), *mexT* and *parR-S* (for MexEF-OprN), and *algU* (for MexCD-OprJ). Since MexAB-OprM can pump out ciprofloxacin we also found duplications in *mexZ*, so as to reduce the production of the competing pump MexXY.

In a combination treatment with all three antibiotics, we observe a similar mutational spectrum as with the ciprofloxacin treatment, where multiple pumps are involved. We performed hierarchical clustering (FigureS2) of these mutated genomes to determine the mutational diversity following each treatment. This figure indicates that the cells have only a few ways to become resistant under meropenem and tobramycin treatment, whereas, they can become resistant to ciprofloxacin through a variety of different mutations. Also, the mutational diversity is lower in Case B vs Case A, when the MexAB pump has a higher expression level compared to the other pumps initially.

### 2.3 Collateral sensitivity and crossresistance

Using the mutated genomes from the three antibiotic treatments, we then assessed whether there were instances of multi-trait phenotype evolution i.e. mutants evolved under one antibiotic having increased/decreased susceptibility to other antibiotics. This is commonly known as collateral sensitivity or cross-resistance respectively. For this analysis, we considered six more antibiotics: colistin, tetracycline, macrolide, trimethoprim, azithromycin and tigecycline that can be extruded by one or several of the four efflux pumps. As a point of comparison, we also carried out mutation simulations in the absence of selection for antibiotic resistance, i.e. where selection only acts to constrain the cost of protein. We considered two constraints on protein costs for evolution in absence of antibiotics: 1. weak selection where the maximum amount of protein must not exceed twice the initial amount and 2. strong selection where the amount of protein must always decrease after every mutation. Figure-3(a) shows that under weak selection for protein cost, the mutated genomes are about equally likely to become susceptible and resistant to the 9 antibiotics. However, under strong selection, the mutated genomes become susceptible to a majority of the antibiotics. This is caused by the fact that each pump consists of multiple genes that contribute more to the protein production cost. Therefore, under strong selection, mutations that lower the expression of all pumps get preferentially selected, increasing the susceptibility of the cells to antibiotics. The notable exception here is the resistance to azithromycin and tigecycline, for the mutants generated in Case B. Due to the higher expression of the MexAB pump in Case B, a common mutation is deletion of *vqsM*, which causes upregulation of the MexCD pump that pumps out these antibiotics.

**Figure 3.**
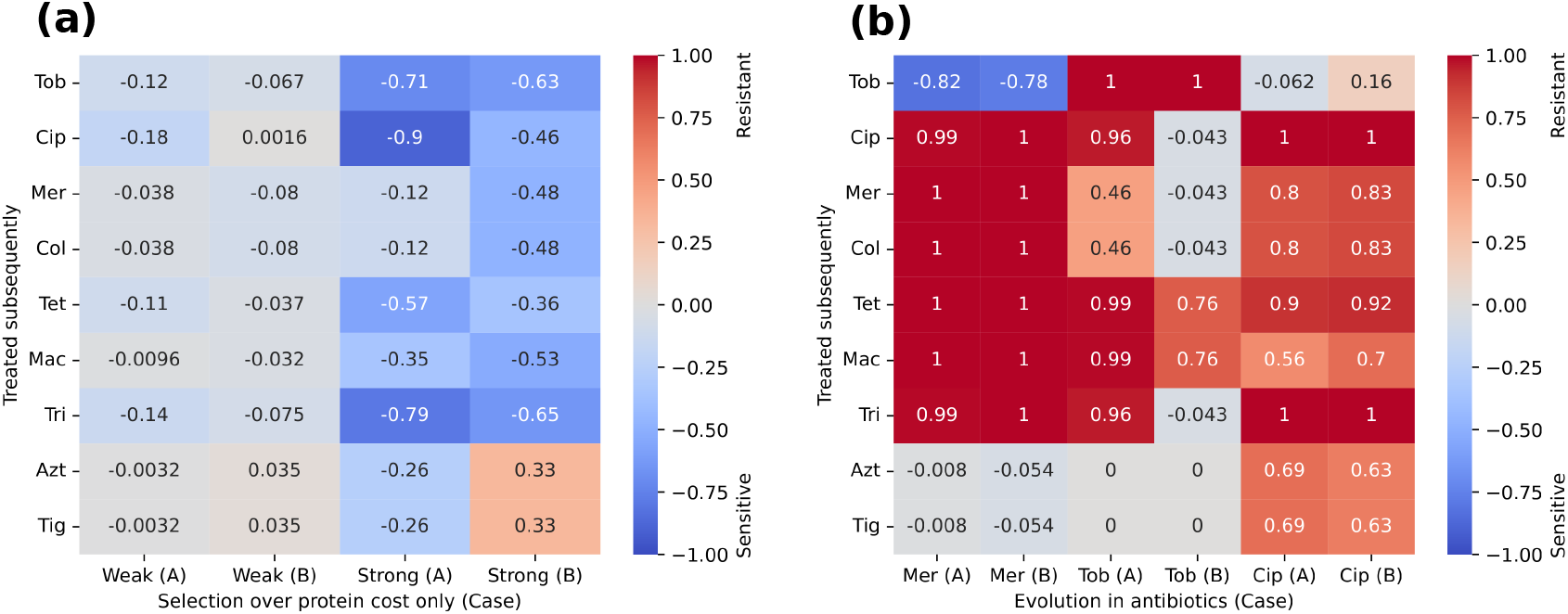
Evolution of cross-resistance and collateral sensitivity between different antibiotics. **(a):** This heatmap shows the fraction of mutated genomes that become sensitive (blue) or resistant (red) to 9 different antibiotics, when evolved without antibiotics. It also shows conditions of weak and strong selection over protein production costs. Evolution without antibiotics leads to bacteria becoming sensitive to most antibiotics, more so under strong selection over protein production cost. **(b):** This heatmap shows the fraction of mutated genomes that become sensitive (blue) or resistant (red) to different antibiotics, when evolved in 3 different antibiotics. Adaptive evolution under antibiotic treatments (when protein cost becomes less important) leads to emergence of cross-resistance between different antibiotics for most cases. A notable case of collateral sensitivity is the enhanced susceptibility to tobramycin following meropenem treatment. In these heatmaps Case (A) and Case (B) denote the two cases of pump expression levels considered.

Next, we considered the possibility of cross-resistance and collateral sensitivity in genomes that evolved under selection for antibiotic resistance. We used the mutated genomes from the tobramycin, meropenem, and ciprofloxacin treatments and checked their susceptibility against all nine antibiotics, i.e. the three used for selection plus six additional ones. Figure-3(b) shows that we find cross-resistance in all treatments and cases. For example, selection for resistance to meropenem resulted in cross-resistance to five other antibiotics. This increased propensity for cross-resistance is caused by a combination of factors: 1. regulators are shared between different pumps; 2. antibiotics can be targeted by multiple pumps; and 3. the same pump can act on multiple antibiotics.

We also found one case in which collateral sensitivity was prominent. The majority (around 80%) of genomes mutated under meropenem become susceptible to tobramycin. This collateral sensitivity is another consequence of the competition between the pumps MexAB and MexXY for OprM. Evolution of meropenem resistance often results in reduced expression of the MexXY pump, which makes the mutated genomes susceptible to tobramycin.

There is also a notable difference in sensitivity for the genomes evolved in tobramycin, between Case A (equal expression of all pumps) and Case B (higher expression of MexAB). For Case A, duplications of *parR-S* and *brlR* are common in tobramycin resistant mutants, which also upregulate the pumps MexAB-OprM and MexEF-OprN.

These duplications causes cross-resistance to other antibiotics that are pumped out by those pumps. Since these duplications are not commonly found in Case B, there is less cross-resistance.

### 2.4 Reversion of resistance

In our simulations cells evolve antibiotic resistance by increasing the expression levels of efflux pump genes. While this is beneficial in the presence of antibiotics, this imposes a cost once antibiotics are removed. Besides the cost of making pump proteins, it also costs energy to keep pumps running— though this running cost is not included in our model. Therefore, we were interested in whether mutants could evolve to the original level of efflux pump expression once antibiotics were removed from the system. We selected the mutants with the highest resistance to antibiotics— those paying the highest protein cost— from each initial set of estimated rates of regulation. We then carried out evolution simulations in which mutations were kept provided that they reduced the amount of pump proteins. This selection criteria is similar to the strong selection from Figure-3(b) except we disregarded the costs of regulatory proteins.

We observed that cells can evolve to lower efflux pump levels, but only in Case A could they return to the original levels prior to antibiotic exposure (see Figure-S3). Figure-4 shows the frequencies of different mutations that led to reduced pump expression. A comparison with Figure-2 indicates that reducing the cost of pumps in the absence of antibiotics requires duplication of regulatory genes (negative regulators) at a higher frequency. But, as evident in Figure-S3, even these mutations are not enough to reach the original pump levels for Case B. This happens in Case B and not Case A because the negative regulators of MexXY, MexCD and MexEF are stronger. Once the cells lose those regulators via mutations, there is no way to get them back, and hence it is difficult to return to the original pump expression levels. These results demonstrate that in order to stop paying the extra costs of efflux pumps in the absence of antibiotics, it is easier to have deleterious mutations in the efflux pumps genes themselves rather than the regulators. We verified this when we allowed mutations in the efflux pump genes of the mutants (see Figure-4).

**Figure 4.**
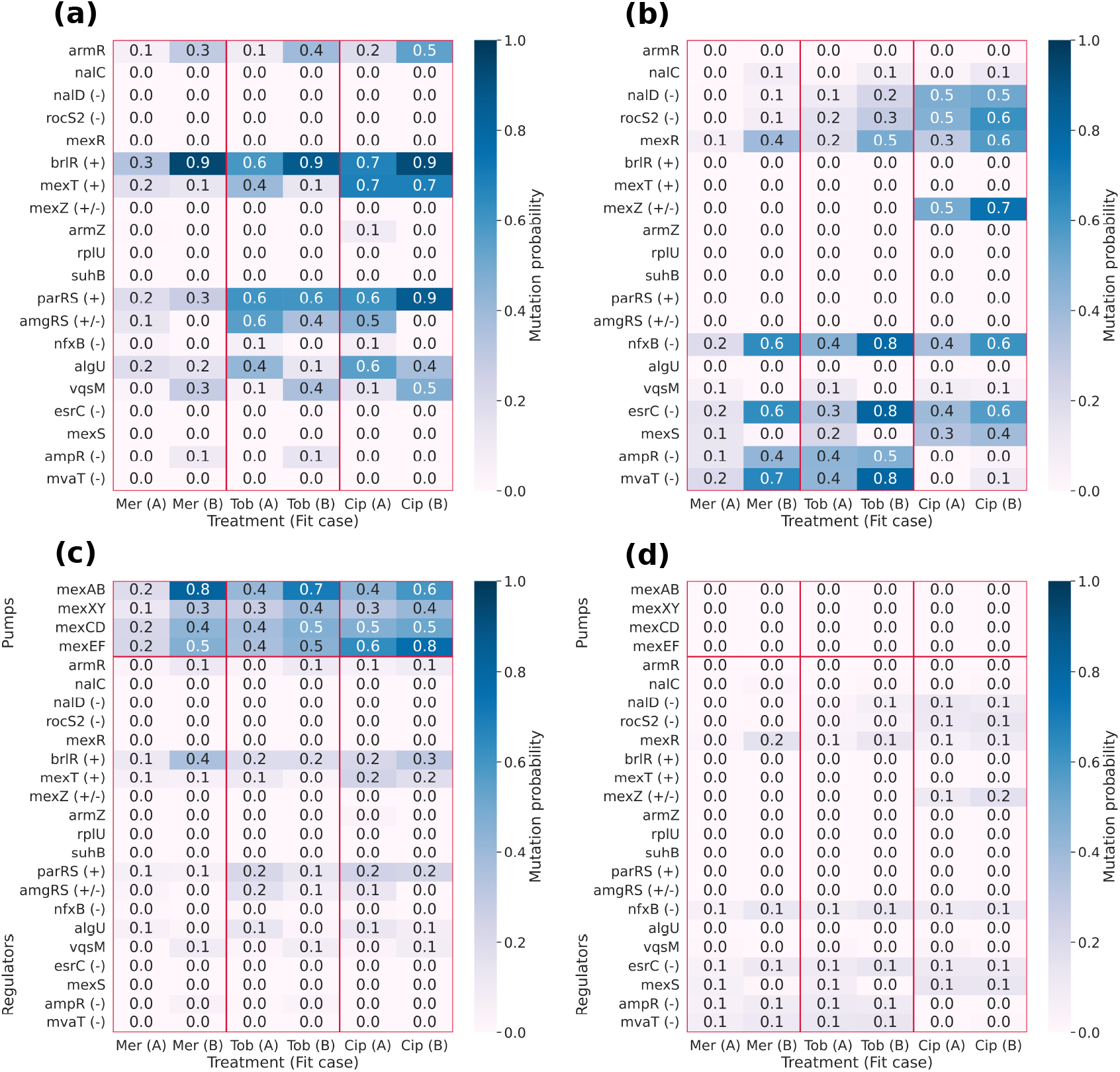
Mutation probabilities for resistant mutants when the antibiotics are removed. **(a)** and **(b):** These heatmaps show the deletion and duplication probabilities respectively for the regulatory genes, when the fittest mutants are evolved without antibiotics. The (+), (-), and (+/-) denote genes that have been previously duplicated, deleted, or both duplicated and deleted depending on the antibiotic used. Reversion of resistance requires more duplications, compared to gaining resistance (see Figure 2(b)). **(c)** and **(d):** These heatmaps show the deletion and duplication probabilities respectively for both efflux pump and regulatory genes, when the mutants are evolved without antibiotics, while allowing for mutations in the efflux pumps genes as well. When the antibiotics are no longer present and the pump-genes are also allowed to mutate, the most likely scenario is deletion of the pump-genes. Case (A) and Case (B) denote two different cases of estimated rates of regulation.

### 2.5 Adapting the model to new conditions

*Pseudomonas aeruginosa* strains have differences in the genomic content of efflux components and regulators, as well as which genes are expressed— or proteins active— in particular environments. For example, the BrlR regulator is a c-di-GMP responsive transcriptional regulator that is active only in cells with high levels of c-di-GMP, like those in biofilms, which is the context modelled in this paper. To simulate evolution in the planktonic state, we removed the *brlR* gene from the network. Moreover, we can include experimental data from in vitro laboratory experiments to better fit parameter values. One example of this is *mexR* which is frequently mutated in planktonic experiments when adapting to ciprofloxacin or meropenem [29, 30] In our modeling of the biofilm context *mexR* rarely gets mutated, because the parameter fitting often leads to it being a weak repressor of the MexAB-OprM pump compared to other regulators such as *nalD* and *rocS2-A2*. We can take into account the experimental data and constrain the parameter fitting to cases in which MexR is a stronger regulator. After doing this, we simulated the planktonic state for Case B, where MexAB has a higher expression compared to other pumps. Figure-5 shows that these modifications led to an increased frequency of *mexR* mutations.

**Figure 5.**
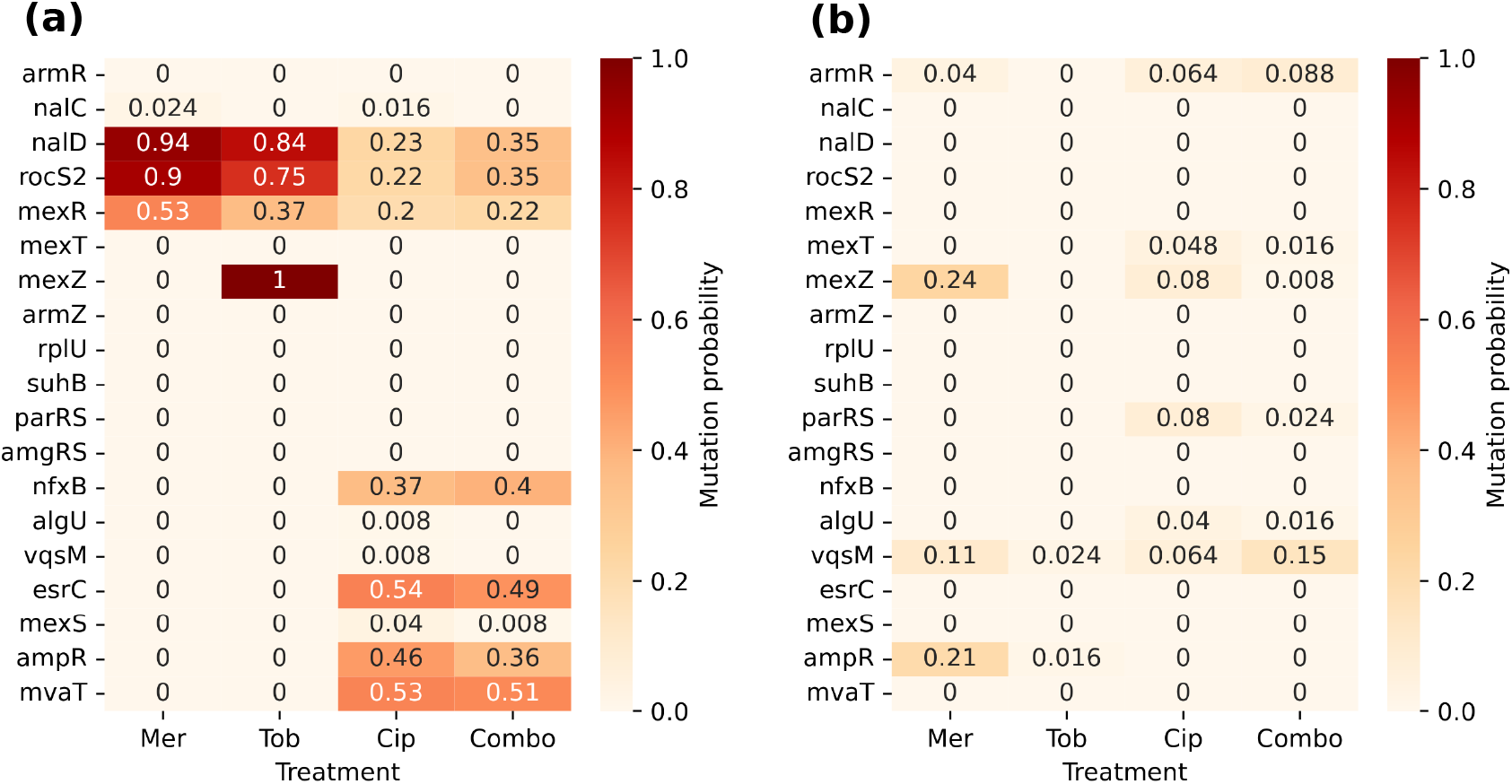
Mutation probabilities of different regulatory genes in the planktonic state. **(a):** This heatmap shows the probabilities of deleterious mutations of the regulatory genes, under different antibiotic treatments. Making *mexR* a strong regulator in the planktonic state increases its deletion probability significantly, compared to the biofilm state. **(b):** This heatmap shows the probabilities of gene duplication mutations of the regulatory genes, when evolved in different antibiotics. The duplication probability of *mexZ* is significantly reduced in the planktonic state. The planktonic state is only simulated for Case B, when the MexAB pump has higher expression compared to the other pumps.

Another notable difference of evolution in the planktonic states compared to biofilms is the rarity of duplications. The gene *mexZ* was a frequent target of duplication in the biofilm state when exposed to meropenem, found in 95-97% of simulations. Its frequency is reduced to less than 25% in the planktonic state. As a result of this, there are fewer instances of collateral sensitivity arising from the MexXY pump (45% and 55% of cases are cross resistant and collaterally sensitive respectively).

## 3 Discussion

The ability to forecast and steer short-term evolution is of fundamental biological interest and has many potential applications especially in infectious diseases and cancer [31]. In addition to the inherent stochasticity of mutations, the complexity of gene regulatory networks often results in mutations having epistastic and pleiotropic effects that make predictions of evolutionary trajectories infeasible. Forecasting the evolution of antibiotic resistance through *de novo* mutations is a particularly suitable approach. Its advantages include the ability to fine-tune selection pressure via antibiotic concentration, well-studied resistance mechanisms, and partly characterized multi-trait effects—where resistance to one antibiotic often increases or decreases resistance to others. Here, we modelled the gene regulatory networks of four efflux pumps and carried out simulations of mutational patterns in the presence and absence of antibiotics. This allowed us to explore the vast combinatorial space of mutations in a complex network and predict cross-resistance and collateral sensitivity after evolution in the presence of three clinically relevant antibiotics. We showed that under ciprofloxacin treatment, the bacteria can become resistant in many different ways while for tobramycin and meropenem mutations were targeted to specific regulatory genes. In all treatments we found repeated patterns: deletions of direct negative regulators and duplications of direct positive regulators. The mutants were multi-trait in the sense that each mutant had reduced/enhanced susceptibility to multiple antibiotics and differing protein costs. So, based on the mutated genomes under these three different antibiotic treatments, we identified the emergence of cross-resistance and collateral sensitivity between these and six more antibiotics. We also showed that when antibiotics are removed, mutants acquire deletion mutations in efflux pump genes to reduce protein expression.

Our model can serve as a null framework for rapidly generating testable forecasts, even when experimental data on regulatory rates or protein concentrations are lacking. One way of evaluating the accuracy of these forecasts is by comparing them with available mutational data from clinical settings or in vitro experiments. Since *P. aeruginosa* often evolves multidrug resistance during chronic lung infections in people with cystic fibrosis, genomic data have been collected on mutations arising under prolonged treatment with multiple antibiotics [32]. From these data we see similar mutational targets as predicted by our model. For example, *mexZ* is one of the most commonly mutated genes in the clinic [32], and it occurs in over 95% of our simulations in response to meropenem and tobramycin. Many additional genes in our model are also found to be mutated multiple times in the clinic, including *nfxB, mexT, mexS, parS, nalC, mexR, nalD, rocS2*, and promoter mutations upstream of *esrC, rplU*. We can also make comparisons with a wealth of *in vitro* data, owing to *P. aeruginosa* being a model organism for antibiotic resistance. Similar to our simulations, adaptive *i*n vitro evolution to ciprofloxacin results in mutations of genes encoding negative regulators including *mexR,nalC, nalD, nfxB* and *mexS* [33, 34]. *Resistance mutations in response to meropenem have been found in nalC, nalD* and *mexR* which are all negative regulators of *mexAB*-*oprM* as well as in *parS* [34, 35]. Also similar to our results, when exposed to a combination of antibiotics, mutations are commonly found in *mexR, nalC*, or *nalD* that all encode negative regulators of *mexAB*-*oprM* and *parS*/*parR* encoding a positive regulator of *mexXY* and *mexEF*-*oprN* [29].

There are also cases where our results differ from *in vitro* or *in vivo* observations. For instance, *in vitro* evolution in response to tobramycin is dominated by mutations in *fusA1*, or genes related to lipopolysaccharides synthesis/modification and the electron transport chain [36, 37]. Since these processes are not included in our regulatory network model, we were unable to identify any associated mutational targets. There are also cases in which genes present in our modeling framework were less often identified as mutational targets than in empirical settings. For example, *algU* appears rarely (*<* 15%) in our simulations in response to ciprofloxacin. Yet, in empirical settings it is a common mutational target. The reason for the discrepancy may be because *algU* encodes a sigma factor and mutating it can modify alginate production and various stress responses [38, 39]. Our modeling framework also over predicted some mutational targets, e.g. mutations in *ampR, mvaT, rocS2* are not commonly found in *in vitro* experiments and mutations in *ampR* and *vqsM* are not found in *in vivo* data. These differences could be due to the role of some of these targets as global regulators, which could cause mutations to have negative epistatic effects.

Our modeling framework can also be used to predict interactions between antibiotics, such as cross-resistance. In our simulations, evolution in the presence of one antibiotic frequently resulted in cross-resistance to multiple other antibiotics, similar to in vitro studies [29, 36, 37, 40, 41]. In some sense this could be predicted in the absence of any simulations, simply based on the fact that efflux pumps can often act on multiple antibiotics. Thus, increasing the expression of a pump will likely lead to an increased ability to remove several antibiotics. However, modeling and simulations can reveal the role of expression levels in shaping cross-resistance patterns. For example, we found that cross-resistance can differ depending on the relative expression level of efflux pumps. When all pumps were equally expressed (Case A) cross-resistance after tobramycin exposure was much more frequent than when *mexAB*-*oprM* had higher relative expression (Case B). Thus, even with the same efflux pumps and the same regulatory network architecture, cross-resistance may depend on the actual expression levels of pumps. These results indicate a benefit in explicit forecasting models that can be tuned to incorporate gene expression information.

Another type of interaction between antibiotics is collateral sensitivity. We found very few instances of collateral sensitivity in our model. It mainly occurred after evolution in response to meropenem, causing increased susceptibility to tobramycin. Similar instances of collateral sensitivity have also been reported in *in vitro* studies where mutants with increased resistance to betalactams (including carbapenems like meropenem) are more susceptible to aminoglycosides like tobramycin [40, 42]. In our model, collateral sensitivity occurs due to downregulation of *mexXY*, which increases availability of the shared OprM protein for MexAB-OprM— the only pump that removes meropenem. There are other examples of collateral sensitivity from empirical observations that our modeling framework did not identify. For instance, *in vitro* studies have found increased susceptibility to aztreonam and colistin after ciprofloxacin resistance evolution [43] and increased susceptibility to meropenem and tobramycin in *nfxB* mutants [44]. This mismatch between our model predictions and empirical data suggests that important regulatory connections may be missing, but once identified they can be included in our model to refine future forecasts.

The fitness of resistant mutants in the absence of antibiotics is a key trait that determines their ability to spread in the environment and between patients in competition with susceptible strains. While fitness costs are common they can also quickly be reduced by either reverting mutations or by compensatory mutations. Importantly, in cases where deletions have resulted in loss of negative regulators, the mutations cannot be directly reverted and proper regulation is lost resulting in high constitutive production of costly efflux pumps. Compensating for this disregulation could then result in further degradation of regulatory networks and efflux pumps. This process could potentially be harnessed to guide bacterial populations into evolutionary dead ends, where they have reduced ability to re-evolve resistance or low fitness due to loss of key genes. We used our model to further explore these questions. After *in silico* evolution in the presence of antibiotics we find, as expected, that production of efflux pumps is increased, which leads to increased protein production costs. These costs remain when antibiotics is removed, and we explored how resistant mutants can evolve to reduce these costs. When allowing only mutations in regulators we find a pattern in deletion of positive regulators and duplication of negative regulators. However, in many cases negative regulators, like *nalD* for meropenem and tobramycin, were already deleted when evolving resistance thereby preventing future mutations and restricting the possible evolutionary trajectories. In the case when we also allow mutation in the genes encoding the efflux pumps, we find that they are commonly deleted, indicating a possibility to guide evolution towards genotypes that do not contain the efflux pump genes, and hence will lack the ability to re-acquire resistance by regulatory mutations. Deletion of efflux pumps genes has been observed in *in vitro* experiments both in the absence of antibiotics [45] or with antibiotics [35, 41, 46] where collateral sensitivity can also select for loss. Interestingly, mutations in efflux pump genes are also among the most commonly mutated genes during in-patient evolution [32] (Table-S1) and deletion of efflux pump genes has even been observed during short-term treatment [47]. In cases where these mutations are irreversible, i.e. large deletions, the reduced potential for the bacteria to re-evolve resistance could be used to inform design of treatment protocols.

Our modeling framework demonstrates the extent to which evolutionary adaptations can be predicted even in the absence of complete empirical data. In developing our model, we necessarily adopted a number of simplifying assumptions, e.g. constant transcription and translation rates across genes as well as equal rates of mutations across genes. Yet, despite these simplifications, the model successfully reproduced evolutionary patterns observed both *in vitro* and in clinical settings. An important part of our approach was the use of genetic algorithms and stochastic evolutionary simulations to explore an ensemble of parameterized networks. By aggregating results across many versions of a network, it reduced the sensitivity to specific parameter choices and instead revealed the predictive power of regulatory structure. The availability of additional information— e.g. expression level differences between planktonic and biofilm growth— could also be incorporated to refine the scope of predictions. The flexibility of this modeling framework allows for the exploration of other possibilities, such as the effects of applying different antibiotics in various sequences. Predictions of the model can then be used to focus experiments on gaining information with predictive power. Repeated iterations of modeling predictions and empirical validations/observations could harness the patterns of antibiotic resistance to create effective evolutionarily-designed treatments.

## 4 Methods

### 4.1 Protein concentrations inside cells

Gene expression can be modeled as a 2-step dynamical process [74] with variables describing the concentrations of mRNA (M) and protein (P):

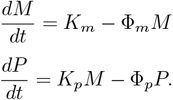

Here *K*_*m*_ and *K*_*p*_ are the transcription and translation rates, and Φ_*m*_ and Φ_*p*_ are the decay rates for mRNA and proteins, respectively. The equations give the following steady values for mRNA and protein:

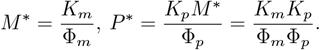

For *K*_*p*_ >> *K*_*m*_, the 2-step process can effectively be considered as a 1-step gene-to-protein creation process with a rate *K*_*tr*_ = *K*_*p*_*K*_*m*_*/*Φ_*m*_ and steady-state protein level *P* ^∗^ = *K*_*tr*_*/*Φ_*p*_. Thus, if we consider a set of different proteins *P*_*i*_ (where *i* is an index) then their abundance can be represented by the set of dynamical equations:

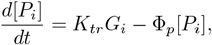

where *G*_*i*_ is the copy number of gene *i*.

Applying this model formulation to *P. aeruginosa*, we can estimate the values of the various rates. The average mRNA and protein levels (per gene) in a *P. aeruginosa* cell are approximately *M* ^∗^ = 0.01 and *P* ^∗^ = 200 [75]. The decline in protein levels is mainly caused by dilution during cell division [76]. Since *Pseudomonas aeruginosa* has a doubling time of 2.3 hours (*h*) [77], the mean rate of protein decay is approximately Φ_*p*_ = *ln*(2)*/*2.3 ∼ 0.3 *h*^−1^ and the rate of gene to protein creation can be estimated as *K*_*tr*_ = *P* ^∗^Φ_*p*_ = 60 *h*^−1^.

To model the action of regulatory proteins that bind to promoter regions and either block or catalyze transcription, the equation is modified as [78]:

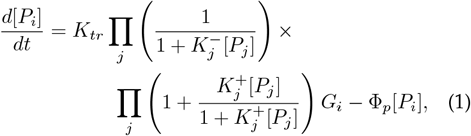

where *K*^+^ and *K*^−^ indicates the rate of regulation of gene *i* by protein *j* (+ sign is for positive regulators and − sign is for negative regulators).

### 4.2 Fitness costs

Fitness in our model system depends on two costs experienced by cells: the cost of protein production and the cost of antibiotics. To estimate both of these costs, we assume that the dynamics of the system in Equation1 are sufficiently fast that we can use the steady state concentrations of proteins. For a given antibiotic, we group together all pumps that target it and call the sum of their steady state values *P*_pump_. Since, each efflux pump consists of 3 proteins and they are transcribed together, the number of efflux pumps (*P*_*pump*_) in a cell can be considered to be equal to the number of either mex (*P*_*mex*_) or opr (*P*_*opr*_) proteins. The exception is the MexXY pump with 2 components which needs OprM sharing from the MexAB-OprM pump. So, we modeled the number of MexAB and MexXY pumps as,

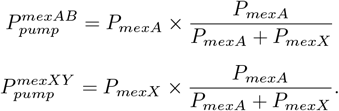

We group the remaining proteins together and call the sum of their steady state values *P*_rest_. The cost of protein production (*C*_*P*_) is then:

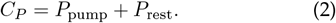

The cost of antibiotic (*C*_*A*_) may depend on many factors including the duration exposed to antibiotics and the efficacy of the pumps. For simplicity, we assume that the cost of antibiotics is inversely proportional to the specific pump(s) that pump out a particular antibiotic 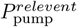, i.e.

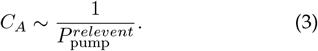

Thus, if there are more pumps the cost of antibiotics will be less because they are actively being removed from cells.

## Supplementary Information

### Robustness of rates of regulation

The rates of regulation *K*^±^ for the gene regulatory network are unknown. Therefore we develop a type of genetic algorithm to fit the initial *K*^±^ network to some desired steady state values of efflux pump proteins such as:

- **A:**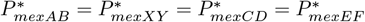
- **B:**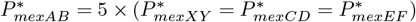

We carried out 25 trials for each case. In each trial, we keep adding random numbers to the *K*^*±*^ values, and only accept the iteration if it reduces the error with respect to desired efflux pump levels. While doing that, we make sure that the regulators only change their strength of regulation and not the their type of regulation (positive and negative). When we compare the difference between each pair of the trials (Figure-S1) we found that each trial generated a different fit. It indicates that networks with different rates of regulation can cause similar expression levels for the efflux pump genes.

---

**Figure S1.**
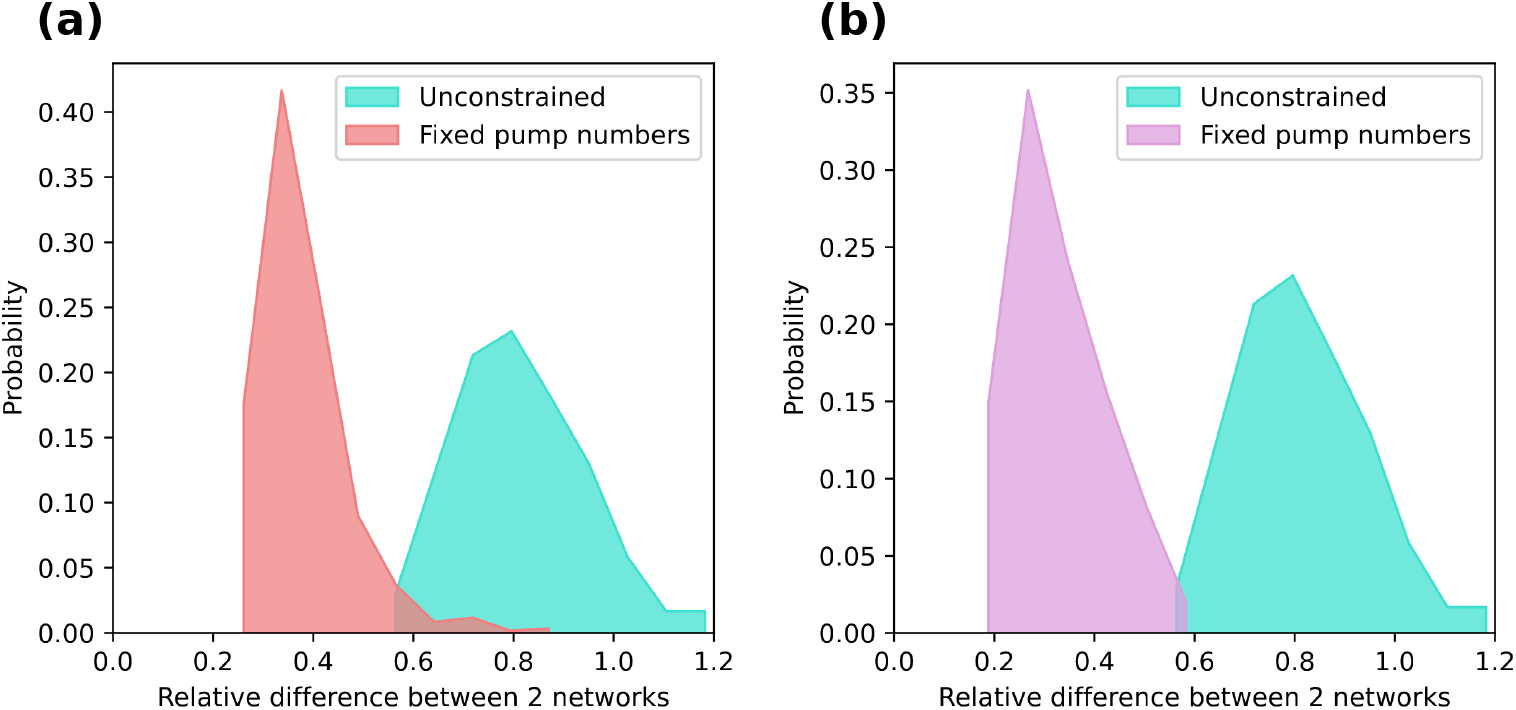
Diversity of the estimated rates of regulation, generated using a genetic algorithm. **(a):** This figure shows the histogram of the relative difference between 2 different sets of estimates, for Case (A) when all 4 pumps have same expression levels. **(b):** This figure shows the histogram of the relative difference between 2 different sets of estimates, for Case (B) when MexAB pump has higher expression level compared to the other pumps. For comparison, each figure also contains a case when the genetic algorithm is applied without any constraint on the expression level of any gene. The figures show that there can moderate level of diversity for the rates of regulation, that can generate a similar expression levels for the efflux pumps.

**Figure S2.**
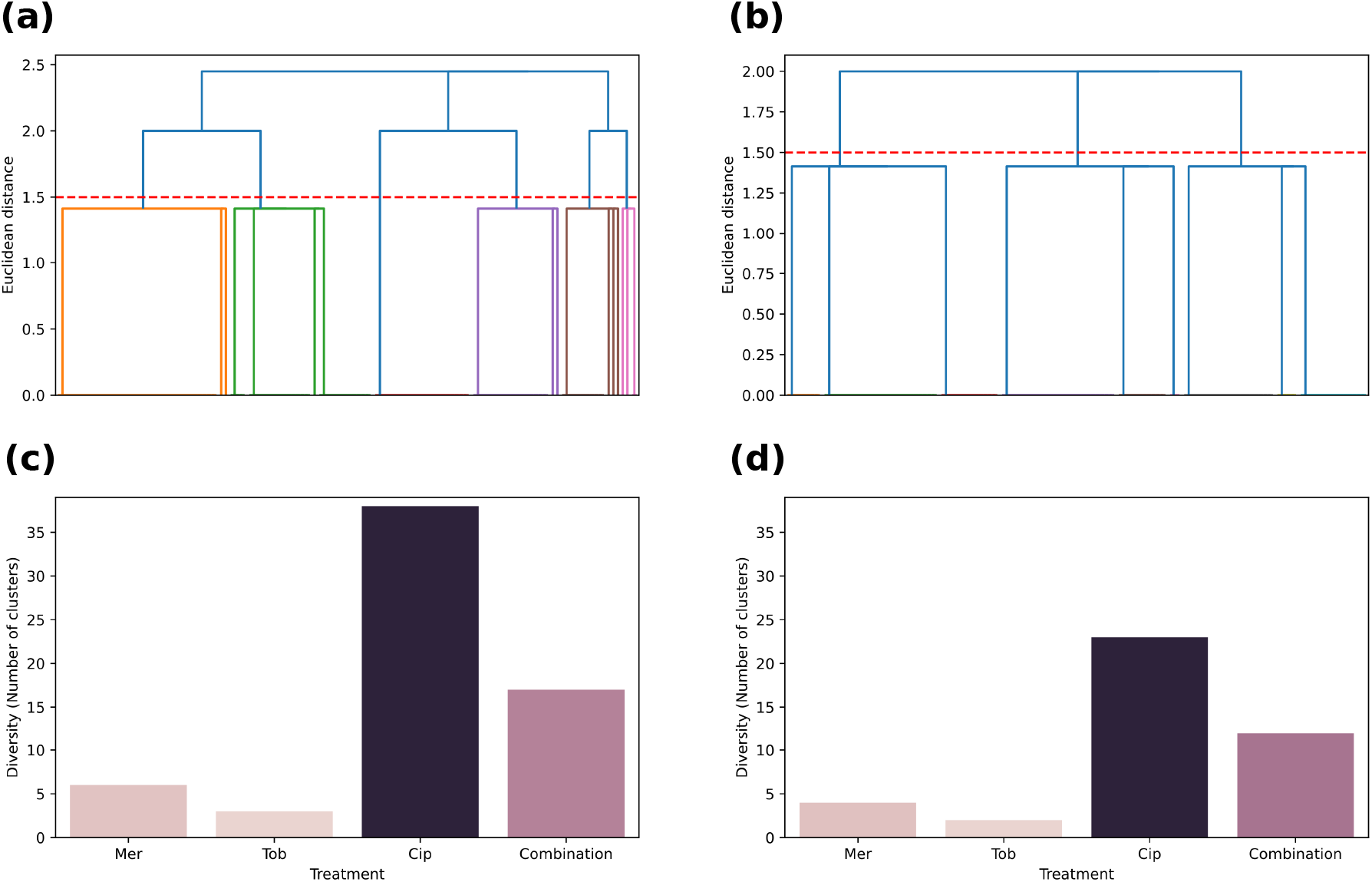
Mutational diversity under different antibiotic treatments. **(a)** and **(b):** These figures show 2 examples of the hierarchical clustering dendrograms, for the mutated genomes following meropenem and tobramycin treatments respectively (in Case A when all pumps have same expression levels). We chose the threshold value to be 1.5 for counting the number of clusters. **(c):** This bar plot shows the diversity of mutated genomes (measured in terms of the number of clusters) for Case A. **(d):** This bar plot shows the diversity of mutated genomes for Case B, when the MexAB pump has higher expression level. The figure shows that evolution under ciprofloxacin treatment can lead to a very high mutational diversity.

**Figure S3.**
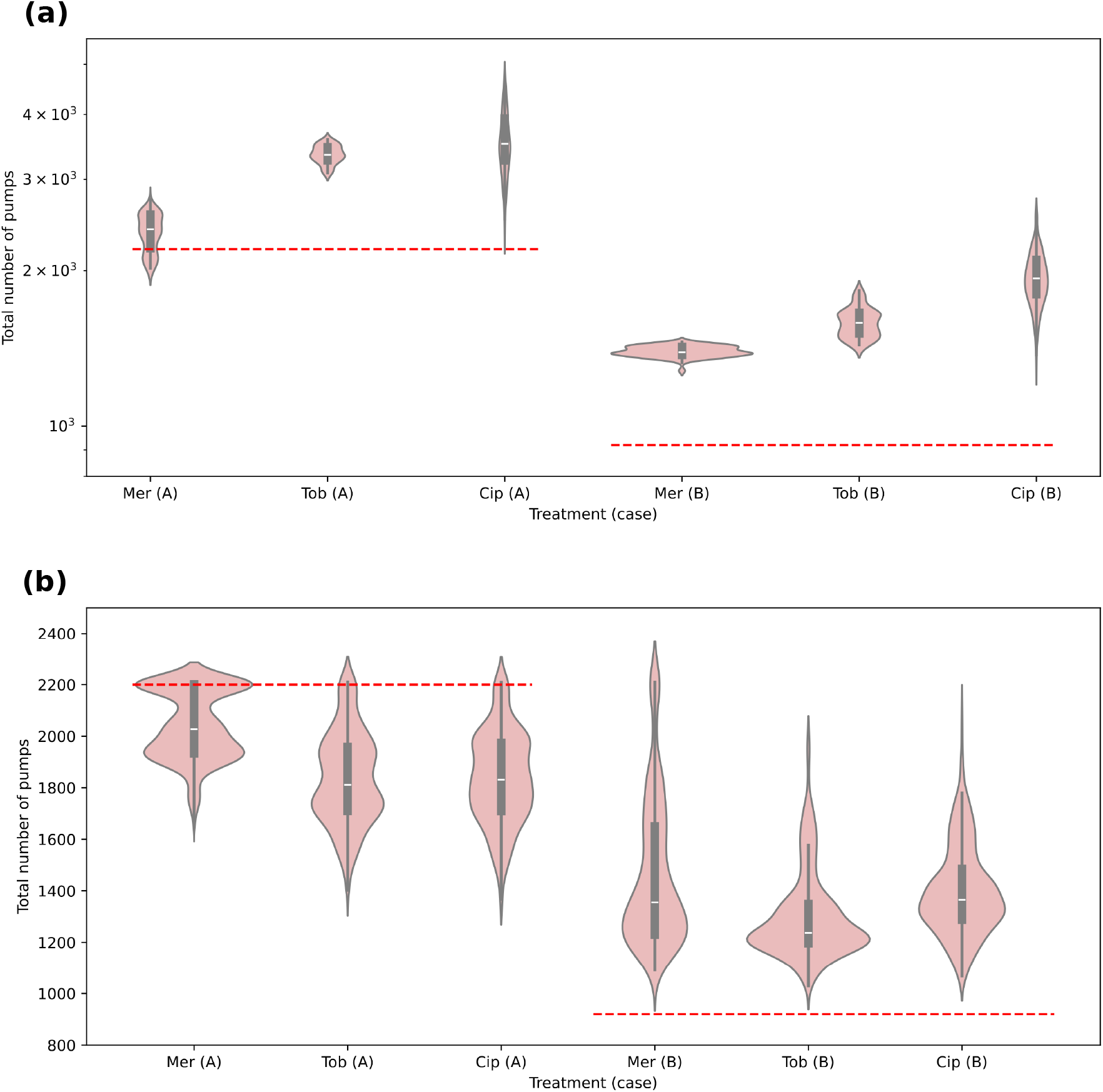
The total number of efflux pumps following different treatments. **(a):** This violin plot shows the number of pumps in the resistant mutants (following treatments) compared to their initial values. Left panel and right panels represent Case A and Case B of initial pump expressions respectively. **(b):** This violin plot shows the number of pumps after the antibiotics are removed and the cells are allow to acquire more mutations in their attempt to go back to the original pump levels. This figure shows that it is not possible to return to the original pump levels in Case B, with mutations in the regulatory genes only.

**Table S1.** Number of unique mutations during within-host evolution in longitudinally collected clinical isolates of *Pseudomonas aeruginosa* sampled from individuals with cystic fibrosis (from [32]).

| Gene | Locus Tag PAO1 | Unique SNPs | Unique Indels | Upstream intergenic |  |
| --- | --- | --- | --- | --- | --- |
| ampR | PA4109 | 0 | 0 | 0 | 0 |
| armR | PA3719 | 0 | 0 | 0 | 0 |
| vqsM | PA2227 | 0 | 0 | 0 | 0 |
| mexX | PA2019 | 1 | 0 | 0 | 1 |
| rocA2 | PA3045 | 1 | 0 | 0 | 1 |
| brlR | PA4878 | 1 | 0 | 0 | 1 |
| suhB | PA3818 | 0 | 1 | 0 | 1 |
| mvaT | PA4315 | 1 | 0 | 0 | 1 |
| oprM | PA0427 | 1 | 1 | 0 | 2 |
| mexY | PA2018 | 2 | 0 | 0 | 2 |
| mexE | PA2493 | 1 | 1 | 0 | 2 |
| armZ | PA5471 | 1 | 1 | 0 | 2 |
| parR | PA1799 | 0 | 0 | 2 | 2 |
| oprN | PA2495 | 3 | 0 | 0 | 3 |
| rocS1 | PA3946 | 3 | 0 | 0 | 3 |
| amgR | PA5200 | 1 | 1 | 1 | 3 |
| oprJ | PA4597 | 4 | 0 | 0 | 4 |
| esrC | PA4596 | 0 | 0 | 4 | 4 |
| rplU | PA4568 | 0 | 0 | 4 | 4 |
| nalD | PA3574 | 0 | 6 | 0 | 6 |
| rocS2 | PA3044 | 3 | 3 | 0 | 6 |
| mexC | PA4599 | 4 | 0 | 3 | 7 |
| mexF | PA2494 | 5 | 2 | 0 | 7 |
| amgS | PA5199 | 6 | 1 | 0 | 7 |
| mexR | PA0424 | 1 | 7 | 0 | 8 |
| nalC | PA3721 | 3 | 4 | 1 | 8 |
| parS | PA1798 | 10 | 0 | 0 | 10 |
| mexS | PA2491 | 9 | 1 | 0 | 10 |
| mexA | PA0425 | 3 | 9 | 0 | 12 |
| mexT | PA2492 | 9 | 5 | 1 | 15 |
| mexD | PA4598 | 17 | 0 | 0 | 17 |
| nfxB | PA4600 | 10 | 8 | 3 | 21 |
| mexB | PA0426 | 6 | 16 | 0 | 22 |
| algU | PA0762 | 22 | 6 | 2 | 30 |
| mexZ | PA2020 | 10 | 30 | 0 | 40 |

## Notes

### Competing Interest Statement

The authors have declared no competing interest.

